# Nanoscale robots exhibiting quorum sensing

**DOI:** 10.1101/448761

**Authors:** Yaniv Amir, Almogit Abu-Horowitz, Justin Werfel, Ido Bachelet

## Abstract

Multi-agent systems demonstrate the ability to collectively perform complex tasks—e.g., construction^1–2^, search^3^, and locomotion^4,5^—with greater speed, efficiency, or effectiveness than could a single agent alone. Direct and indirect coordination methods allow agents to collaborate to share information and adapt their activity to fit dynamic situations. A well-studied example is quorum sensing (QS), a mechanism allowing bacterial communities to coordinate and optimize various phenotypes in response to population density. Here we implement, for the first time, bio-inspired QS in robots fabricated from DNA origami, which communicate by transmitting and receiving diffusing signals. The mechanism we describe includes features such as programmable response thresholds and quorum quenching, and is capable of being triggered by proximity of a specific target cell. Nanoscale robots with swarm intelligence could carry out tasks that have been so far unachievable in diverse fields such as industry, manufacturing and medicine.

## Introduction, Results, and Discussion

Quorum Sensing (QS) is a well-studied example of collective behavior^6^. This mechanism of cell-cell communication in bacteria utilizes secreted signal molecules to coordinate the behavior of the group. Linking signal concentration to local population density enables each single bacterium to measure population size. This ability to communicate both within and between species is critical for bacterial survival and interaction in natural habitats and has likely appeared early in evolution. Detection of a minimal threshold of signal molecules, termed autoinducers, triggers gene expression and subsequent behavior response. Using these signaling systems, bacteria synchronize particular behaviors on a population-wide scale and thus function as multicellular organisms^6–9^.

QS-inspired approaches have been adopted in artificial systems, including mobile robots^10^ and wireless sensor networks^11^, and naturally occurring genes have been harnessed in synthetic biology to implement QS at the cellular level^12^.

Recently we reported a new type of nanoscale robot, fabricated from DNA origami^13^, which logically actuates between “off” and “on” states^14–15^. By using various types of DNA logic based on aptamer recognition, toehold-mediated strand displacement^15^, etc., these robots can be programmed to respond to diverse stimuli and either present or sequester molecular payloads anchored to the inside of the device. In the present study we aimed to program the robots to exhibit collective behavior, taking advantage of the more elaborate modes of control that such behaviors enable.

The basis for collective behaviors is communication between agents, and QS was chosen as a simple, programmable mechanism to establish it. We designed and constructed a bio-inspired QS system based on an autoinducer which is released by each individual robot into the environment **(Supplementary notes 1, 2)**. The inducer is in fact the key that opens the robot, which is initially bound to the internal face of every robot. The quantity of this molecule is thus proportional to the robot population size, and each individual robot is able to detect it and respond in a concentration-dependent fashion. To achieve this, a key previously utilized to open the robot, recombinant human platelet-derived growth factor (PDGF)^14–15^, was used as an autoinducer. PDGF was loaded into the robots using a peptide tether cleavable by matrix metalloproteinase (MMP)-2 **(Fig. 1A, Fig. S1-S3)**. Thus, MMP-2 was set as an external signal initiating QS, and changing its activity enabled us to tune the rate of autoinducer release. Importantly, due to the hollow cylindrical shape of the robot, MMP-2 can freely diffuse in and out of the robot and operate inside it in both its closed and open states, while PDGF has to be cleaved and released from the robot by MMP-2, in order for other robots to sense and respond to it **(Fig. 1B, C)**.

**Figure 1:**
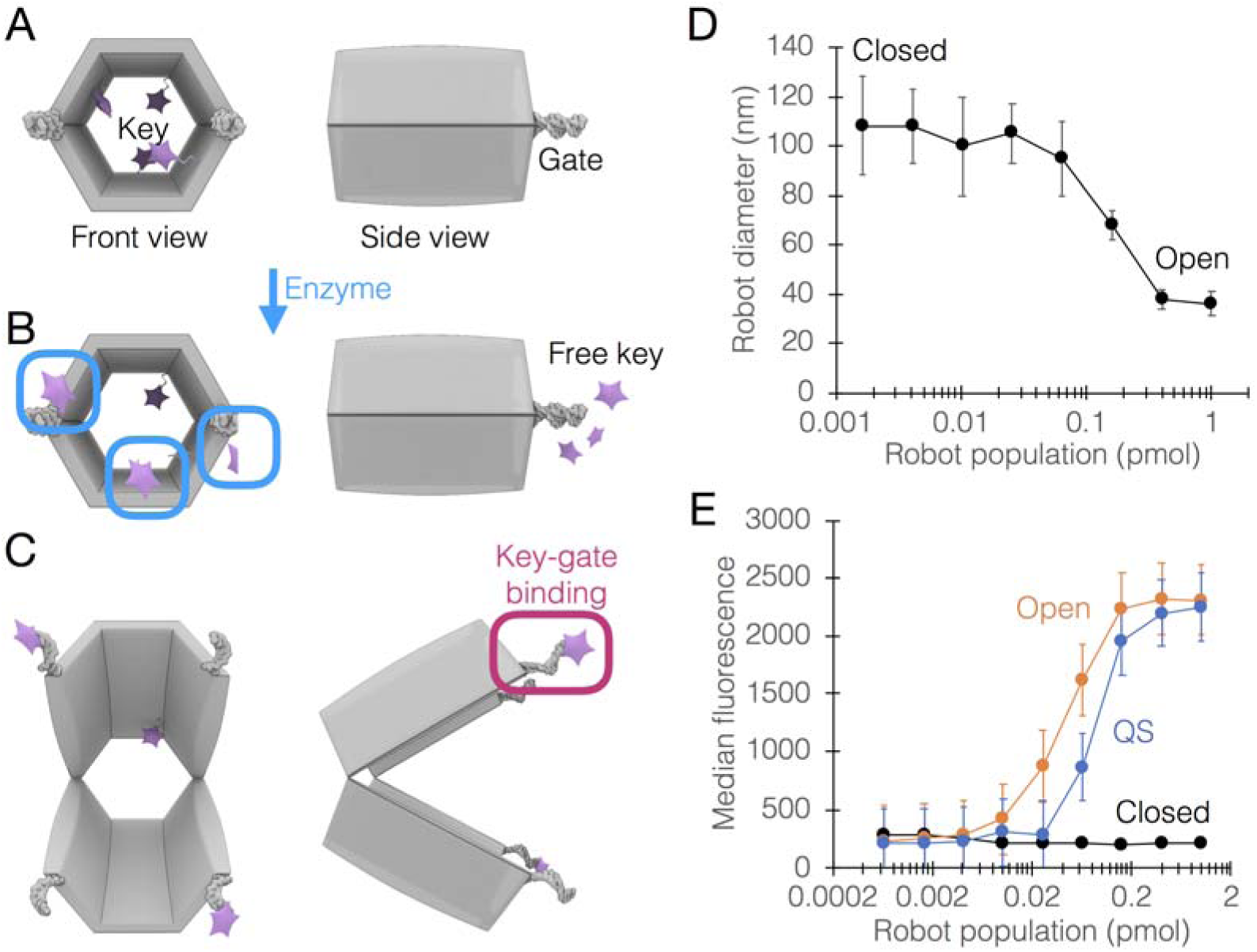
QS in DNA robots. **A-C**, Schematic design of QS system. **A**, The key, platelet-derived growth factor (PDGF, purple stars) was linked chemically to an MMP2-cleavable peptide tether, to form the autoinducer (see **Fig. S1** for zoom-in). A mixture of autoinducer and GFP was loaded inside the robot (top left, seen from the side). **B**, MMP2 releases the autoinducers from a transmitting robot (in a closed state), these reach a receiving robot, switching it from closed to open (top right). **D-E**, Population-dependent behavior of QS robots. Robots were placed in MMP2-containing buffer in various population sizes in a fixed volume and their state was monitored using dynamic light scattering **(D)** or flow cytometry **(E)**, using beads coated with anti-GFP antibodies.

The autoinducer release mechanism can be potentially adapted to any environment. For example, one could exploit the inherent instability of RNA for the gradual release of signal from the robots. Alternatively, a UV-cleavable tether would release the signal only upon exposure of the robots to sunlight or another direct source of UV radiation. Choosing enzymes such as MMPs as releasing factors has a therapeutic rationale, as it only initiates QS where enzyme activity is enriched, such as around or directly on metastasizing tumors^16^.

The closed robots are hollow shells enabling small molecules such as proteins to freely diffuse in and out of them. Specifically here, the protein diffusing in and out is the release factor MMP-2, which when inside releases PDGF (tethered to the robot by the MMP-2 substrate polypeptide). The released PDGF can now also freely diffuse out of the robot, and build up a concentration of PDGF in the environment. In contrast, any attached payload (e.g. reporter molecule or unreleased PDGF) is only accessible to beads or other solid phase-based assays when the robot is open. Therefore, all robots – closed and open – participate in generating the PDGF concentration in the environment, but only the open robots contribute to the detectable signal. Robots loaded with auotoinducers were placed in MMP-2-containing buffer at various population densities (from 29 to 18,000 pM). Population density-dependent activation of the robots was demonstrated using both flow cytometry and dynamic light scattering analysis **(Fig. 1D)**. Flow cytometry clearly showed distinct, QS-driven robot activation behavior displayed between the constitutively off (closed) and constitutively on (open) curves **(Fig. 1E)**.

Engineered QS enables the tuning of response thresholds to fit various conditions or desired behaviors. Here this was achieved by modifying the aptamer gate that responds to the QS signal. In the robot, the aptamer that binds the autoinducer is normally hybridized to a partially-complementary strand, from which it displaces in the presence of the signal as previously shown^14–15^. By changing the number of mismatches in the complementary strand, displacement can be made to occur at lower signal concentrations and with faster kinetics. We used this approach to successfully alter QS-driven behavior in robots **(Fig. 2A, B)**.

**Figure 2:**
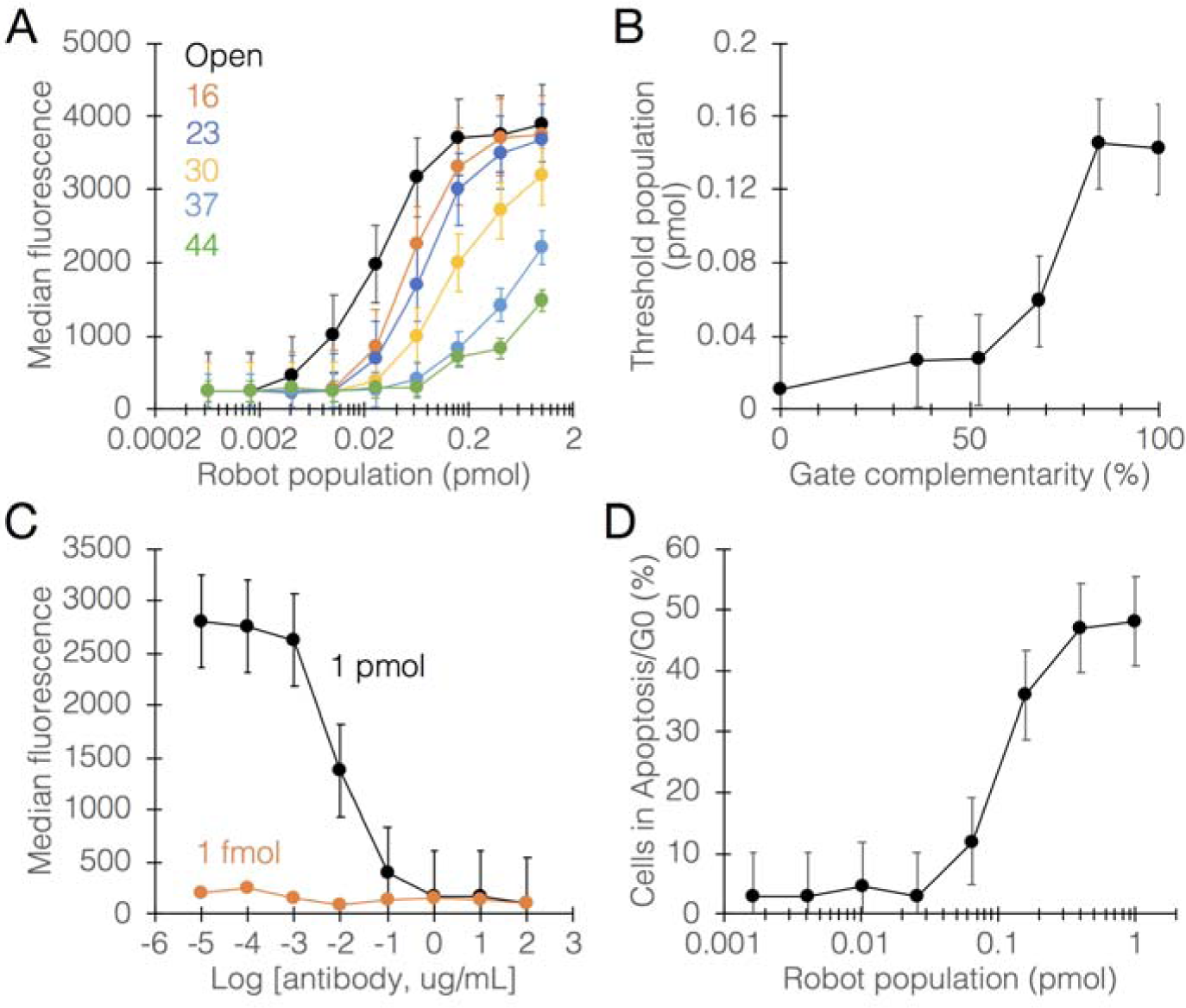
Tuning the behavior of QS robots. QS behavior can be tuned through either the autoinducer-sensing mechanism or through sequestering the autoinducer itself. **A**, reducing complementarity between the DNA strands comprising the robot gate enables to tune the threshold and kinetics of QS behavior. Each curve corresponds to a number of matching bases (max. complementarity: 44 bases; min. complementarity: 16 bases). **B**, quantitative link between % complementarity and the threshold population of robots, i.e. the first population with detectable effect. **C**, sequestration of the autoinducer by a neutralizing anti-PDGF antibody enables quorum quenching (QQ). **D**, Cytokine-activated Jurkat T cells were treated with QS robots loaded with a growth-suppressing antibody (anti-Siglec-7). No MMP-2 was added to the medium as in the previous experiments. QS was driven by MMP2 released from the target cells, leading to subsequent growth arrest. Cells were fixed after treatment and analyzed for cell cycle distribution by flow cytometry.

Our QS system can be tuned also via quorum quenching (QQ), by neutralizing or sequestering the autoinducer. To achieve QQ, we used a neutralizing anti-PDGF antibody^14^ that effectively negated PDGF binding to its aptamer on the robot, causing robots to switch to off even though their concentration was high enough to induce QS-driven activation **(Fig. 2C)**. The efficacy of QQ depended on the ability of the neutralizing antibody to compete with the aptamers for autoinducer binding.

We next loaded the robots with antibody Fab’ fragments for the human receptor Siglec-7 (CDw328), whose cross-linking on leukemic cells induces growth arrest leading to apoptosis^17^. Jurkat cells (leukemic T cells) were chosen as target cells as they express Siglec-7^18^ and also exhibit high levels of MMP-2 activity after activation with cytokines^19^. The cells were treated with varying concentrations of QS-regulated robots for 24 hours. Cell cycle analysis demonstrated cell-triggered QS leading to robot activation and subsequent growth arrest, as no other releasing factor was added to the medium **(Fig 2D)**. This highlights the potential of QS as an artificial therapeutic control mechanism that could be utilized in a variety of conditions, given that the proper system is designed with a target-associated releasing factor in mind, such as tumor-derived proteases, bacterial restriction nucleases, etc. A library of autoinducer tethers, each cleavable by a different signal, could be constructed to fit specific needs and environmental conditions.

In this work we implement, for the first time, collective behavior in molecular robots using a bio-inspired mechanism. The design presented here bears many similarities to bacterial QS, while carrying additional features such as the ability to be activated in response to chosen stimuli. Our work also provides a platform for the engineering of more elaborate communication schemes utilizing several sub-populations differing in autoinducer type and response thresholds, with desirable features as control systems for therapeutics and manufacturing.

## Acknowledgements

We are extremely grateful to S. M. Douglas, D. Y. Zhang and A. Marblestone for their valuable advice and comments on the manuscript. We thank all the members of the Bachelet lab at Bar Ilan University for support, technical help and valuable discussions. This work was supported by a European Research Council Starting Grant (ERC-StG-335332).

## Supplementary Figure 1

**Figure S3.**
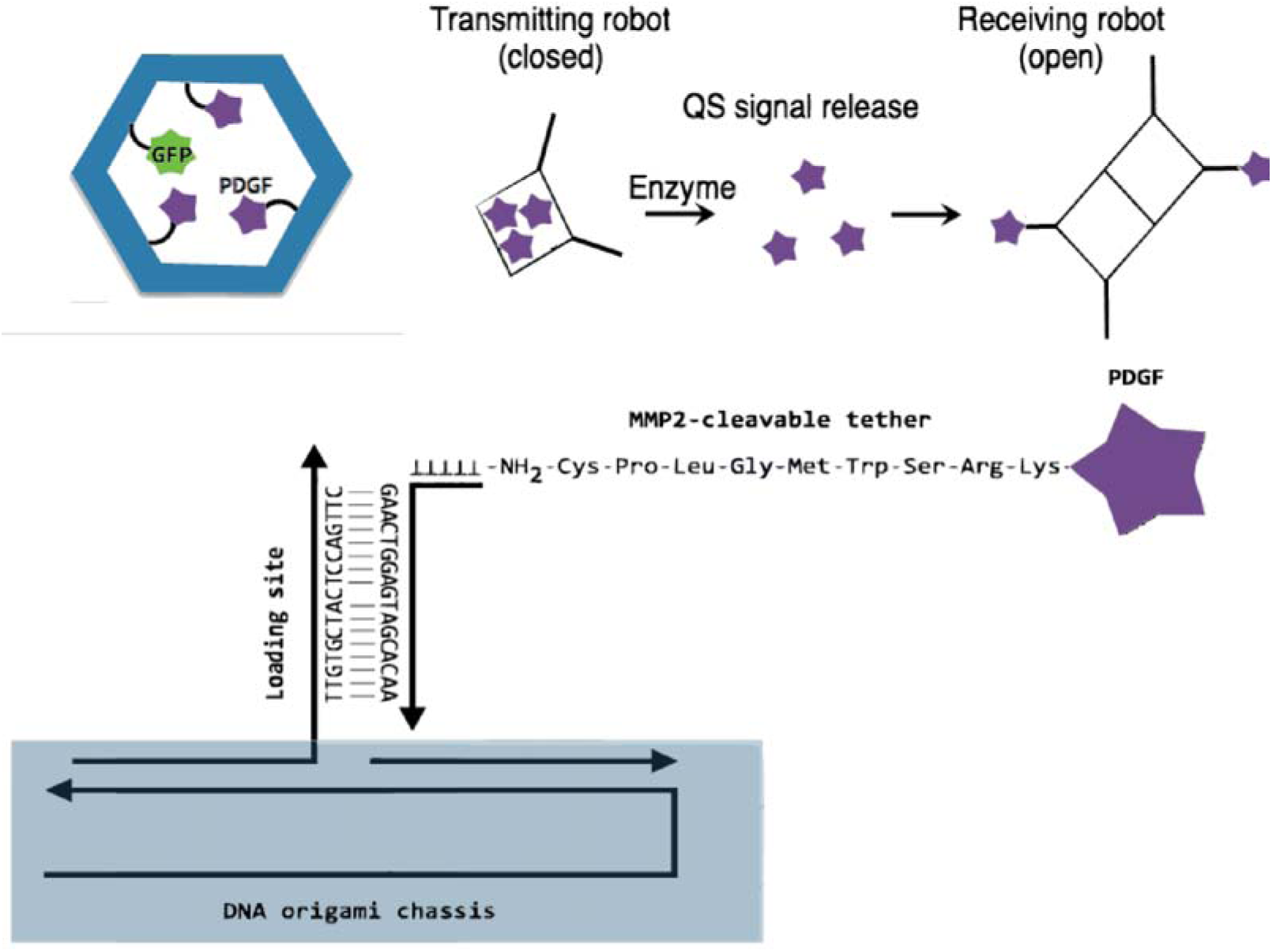
Schematic zoom-in on the linkage between PDGF and robot.

