## Supplementary notes for "Nanoscale robots exhibiting quorum sensing"

### Supplementary Note 1: robot design and fabrication

DNA origami robots were designed using caDNAno (http://www.cadnano.org) and fabricated as previously described. The M13mp18 circular ssDNA was used as scaffold strand. Staple strands were ordered from Integrated DNA technologies.

### Supplementary Table 1: M13mp18 sequence

```
TAGTTGCATATTTAAAACATGTTGAGCTACAGCATTATATTCAGCAATTAAGCTCTAAGCCATCCGCAAAAATGACCTCTTATCAAAAG
GAGCAATTAAAGGTACTCTCTAATCCTGACCTGTTGGAGTTTGCTTCCGGTCTGGTTCGCTTTGAAGCTCGAATTAAAACGCGATATTT
GAAGTCTTTCGGGCTTCCTCTTAATCTTTTTGATGCAATCCGCTTTGCTTCTGACTATAATAGTCAGGGTAAAGACCTGATTTTTGATT
TATGGTCATTCTCGTTTTCTGAACTGTTTAAAGCATTTGAGGGGGATTCAATGAATATTTATGACGATTCCGCAGTATTGGACGCTATC
CAGTCTAAACATTTTACTATTACCCCCTCTGGCAAAACTTCTTTTGCAAAAGCCTCTCGCTATTTTGGTTTTTATCGTCGTCTGGTAAA
CGAGGGTTATGATAGTGTTGCTCTTACTATGCCTCGTAATTCCTTTTGGCGTTATGTATCTGCATTAGTTGAATGTGGTATTCCTAAAT
CTCAACTGATGAATCTTTCTACCTGTAATAATGTTGTTCCGTTAGTTCGTTTTATTAACGTAGATTTTTCTTCCCAACGTCCTGACTGG
TATAATGAGCCAGTTCTTAAAATCGCATAAGGTAATTCACAATGATTAAAGTTGAAATTAAACCATCTCAAGCCCAATTTACTACTCGT
TCTGGTGTTTCTCGTCAGGGCAAGCCTTATTCACTGAATGAGCAGCTTTGTTACGTTGATTTGGGTAATGAATATCCGGTTCTTGTCAA
GATTACTCTTGATGAAGGTCAGCCAGCCTATGCGCCTGGTCTGTACACCGTTCATCTGTCCTCTTTCAAAGTTGGTCAGTTCGGTTCCC
TTATGATTGACCGTCTGCGCCTCGTTCCGGCTAAGTAACATGGAGCAGGTCGCGGATTTCGACACAATTTATCAGGCGATGATACAAAT
CTCCGTTGTACTTTGTTTCGCGCTTGGTATAATCGCTGGGGGTCAAAGATGAGTGTTTTAGTGTATTCTTTTGCCTCTTTCGTTTTAGG
TTGGTGCCTTCGTAGTGGCATTACGTATTTTACCCGTTTAATGGAAACTTCCTCATGAAAAAGTCTTTAGTCCTCAAAGCCTCTGTAGC
CGTTGCTACCCTCGTTCCGATGCTGTCTTTCGCTGCTGAGGGTGACGATCCCGCAAAAGCGGCCTTTAACTCCCTGCAAGCCTCAGCGA
CCGAATATATCGGTTATGCGTGGGCGATGGTTGTTGTCATTGTCGGCGCAACTATCGGTATCAAGCTGTTTAAGAAATTCACCTCGAAA
GCAAGCTGATAAACCGATACAATTAAAGGCTCCTTTTGGAGCCTTTTTTTTGGAGATTTTCAACGTGAAAAAATTATTATTCGCAATTC
CTTTAGTTGTTCCTTTCTATTCTCACTCCGCTGAAACTGTTGAAAGTTGTTTAGCAAAATCCCATACAGAAAATTCATTTACTAACGTC
TGGAAAGACGACAAAACTTTAGATCGTTACGCTAACTATGAGGGCTGTCTGTGGAATGCTACAGGCGTTGTAGTTTGTACTGGTGACGA
AACTCAGTGTTACGGTACATGGGTTCCTATTGGGCTTGCTATCCCTGAAAATGAGGGTGGTGGCTCTGAGGGTGGCGGTTCTGAGGGTG
GCGGTTCTGAGGGTGGCGGTACTAAACCTCCTGAGTACGGTGATACACCTATTCCGGGCTATACTTATATCAACCCTCTCGACGGCACT
TATCCGCCTGGTACTGAGCAAAACCCCGCTAATCCTAATCCTTCTCTTGAGGAGTCTCAGCCTCTTAATACTTTCATGTTTCAGAATAA
TAGGTTCCGAAATAGGCAGGGGGCATTAACTGTTTATACGGGCACTGTTACTCAAGGCACTGACCCCGTTAAAACTTATTACCAGTACA
CTCCTGTATCATCAAAAGCCATGTATGACGCTTACTGGAACGGTAAATTCAGAGACTGCGCTTTCCATTCTGGCTTTAATGAGGATTTA
TTTGTTTGTGAATATCAAGGCCAATCGTCTGACCTGCCTCAACCTCCTGTCAATGCTGGCGGCGGCTCTGGTGGTGGTTCTGGTGGCGG
CTCTGAGGGTGGTGGCTCTGAGGGTGGCGGTTCTGAGGGTGGCGGCTCTGAGGGAGGCGGTTCCGGTGGTGGCTCTGGTTCCGGTGATT
TTGATTATGAAAAGATGGCAAACGCTAATAAGGGGGCTATGACCGAAAATGCCGATGAAAACGCGCTACAGTCTGACGCTAAAGGCAAA
CTTGATTCTGTCGCTACTGATTACGGTGCTGCTATCGATGGTTTCATTGGTGACGTTTCCGGCCTTGCTAATGGTAATGGTGCTACTGG
TGATTTTGCTGGCTCTAATTCCCAAATGGCTCAAGTCGGTGACGGTGATAATTCACCTTTAATGAATAATTTCCGTCAATATTTACCTT
CCCTCCCTCAATCGGTTGAATGTCGCCCTTTTGTCTTTGGCGCTGGTAAACCATATGAATTTTCTATTGATTGTGACAAAATAAACTTA
TTCCGTGGTGTCTTTGCGTTTCTTTTATATGTTGCCACCTTTATGTATGTATTTTCTACGTTTGCTAACATACTGCGTAATAAGGAGTC
TTAATCATGCCAGTTCTTTTGGGTATTCCGTTATTATTGCGTTTCCTCGGTTTCCTTCTGGTAACTTTGTTCGGCTATCTGCTTACTTT
TCTTAAAAAGGGCTTCGGTAAGATAGCTATTGCTATTTCATTGTTTCTTGCTCTTATTATTGGGCTTAACTCAATTCTTGTGGGTTATC
TCTCTGATATTAGCGCTCAATTACCCTCTGACTTTGTTCAGGGTGTTCAGTTAATTCTCCCGTCTAATGCGCTTCCCTGTTTTTATGTT
ATTCTCTCTGTAAAGGCTGCTATTTTCATTTTTGACGTTAAACAAAAAATCGTTTCTTATTTGGATTGGGATAAATAATATGGCTGTTT
ATTTTGTAACTGGCAAATTAGGCTCTGGAAAGACGCTCGTTAGCGTTGGTAAGATTCAGGATAAAATTGTAGCTGGGTGCAAAATAGCA
ACTAATCTTGATTTAAGGCTTCAAAACCTCCCGCAAGTCGGGAGGTTCGCTAAAACGCCTCGCGTTCTTAGAATACCGGATAAGCCTTC
TATATCTGATTTGCTTGCTATTGGGCGCGGTAATGATTCCTACGATGAAAATAAAAACGGCTTGCTTGTTCTCGATGAGTGCGGTACTT
GGTTTAATACCCGTTCTTGGAATGATAAGGAAAGACAGCCGATTATTGATTGGTTTCTACATGCTCGTAAATTAGGATGGGATATTATT
TTTCTTGTTCAGGACTTATCTATTGTTGATAAACAGGCGCGTTCTGCATTAGCTGAACATGTTGTTTATTGTCGTCGTCTGGACAGAAT
TACTTTACCTTTTGTCGGTACTTTATATTCTCTTATTACTGGCTCGAAAATGCCTCTGCCTAAATTACATGTTGGCGTTGTTAAATATG
GCGATTCTCAATTAAGCCCTACTGTTGAGCGTTGGCTTTATACTGGTAAGAATTTGTATAACGCATATGATACTAAACAGGCTTTTTCT
AGTAATTATGATTCCGGTGTTTATTCTTATTTAACGCCTTATTTATCACACGGTCGGTATTTCAAACCATTAAATTTAGGTCAGAAGAT
GAAATTAACTAAAATATATTTGAAAAAGTTTTCTCGCGTTCTTTGTCTTGCGATTGGATTTGCATCAGCATTTACATATAGTTATATAA
CCCAACCTAAGCCGGAGGTTAAAAAGGTAGTCTCTCAGACCTATGATTTTGATAAATTCACTATTGACTCTTCTCAGCGTCTTAATCTA
AGCTATCGCTATGTTTTCAAGGATTCTAAGGGAAAATTAATTAATAGCGACGATTTACAGAAGCAAGGTTATTCACTCACATATATTGA
TTTATGTACTGTTTCCATTAAAAAAGGTAATTCAAATGAAATTGTTAAATGTAATTAATTTTGTTTTCTTGATGTTTGTTTCATCATCT
TCTTTTGCTCAGGTAATTGAAATGAATAATTCGCCTCTGCGCGATTTTGTAACTTGGTATTCAAAGCAATCAGGCGAATCCGTTATTGT
TTCTCCCGATGTAAAAGGTACTGTTACTGTATATTCATCTGACGTTAAACCTGAAAATCTACGCAATTTCTTTATTTCTGTTTTACGTG
CAAATAATTTTGATATGGTAGGTTCTAACCCTTCCATTATTCAGAAGTATAATCCAAACAATCAGGATTATATTGATGAATTGCCATCA
TCTGATAATCAGGAATATGATGATAATTCCGCTCCTTCTGGTGGTTTCTTTGTTCCGCAAAATGATAATGTTACTCAAACTTTTAAAAT
TAATAACGTTCGGGCAAAGGATTTAATACGAGTTGTCGAATTGTTTGTAAAGTCTAATACTTCTAAATCCTCAAATGTATTATCTATTG
ACGGCTCTAATCTATTAGTTGTTAGTGCTCCTAAAGATATTTTAGATAACCTTCCTCAATTCCTTTCAACTGTTGATTTGCCAACTGAC
CAGATATTGATTGAGGGTTTGATATTTGAGGTTCAGCAAGGTGATGCTTTAGATTTTTCATTTGCTGCTGGCTCTCAGCGTGGCACTGT
```

TGCAGGCGGTGTTAATACTGACCGCCTCACCTCTGTTTTATCTTCTGCTGGTGGTTCGTTCGGTATTTTTAATGGCGATGTTTTAGGGC
TATCAGTTCGCGCATTAAAGACTAATAGCCATTCAAAAATATTGTCTGTGCCACGTATTCTTACGCTTTCAGGTCAGAAGGGTTCTATC
TCTGTTGGCCAGAATGTCCCTTTTATTACTGGTCGTGTGACTGGTGAATCTGCCAATGTAAATAATCCATTTCAGACGATTGAGCGTCA
AAATGTAGGTATTTCCATGAGCGTTTTTCCTGTTGCAATGGCTGGCGGTAATATTGTTCTGGATATTACCAGCAAGGCCGATAGTTTGA
GTTCTTCTACTCAGGCAAGTGATGTTATTACTAATCAAAGAAGTATTGCTACAACGGTTAATTTGCGTGATGGACAGACTCTTTTACTC
GGTGGCCTCACTGATTATAAAAACACTTCTCAGGATTCTGGCGTACCGTTCCTGTCTAAAATCCCTTTAATCGGCCTCCTGTTTAGCTC
CCGCTCTGATTCTAACGAGGAAAGCACGTTATACGTGCTCGTCAAAGCAACCATAGTACGCGCCCTGTAGCGGCGCATTAAGCGCGGCG
GGTGTGGTGGTTACGCGCAGCGTGACCGCTACACTTGCCAGCGCCCTAGCGCCCGCTCCTTTCGCTTTCTTCCCTTCCTTTCTCGCCAC
GTTCGCCGGCTTTCCCCGTCAAGCTCTAAATCGGGGGCTCCCTTTAGGGTTCCGATTTAGTGCTTTACGGCACCTCGACCCCAAAAAAC
TTGATTTGGGTGATGGTTCACGTAGTGGGCCATCGCCCTGATAGACGGTTTTTCGCCCTTTGACGTTGGAGTCCACGTTCTTTAATAGT
GGACTCTTGTTCCAAACTGGAACAACACTCAACCCTATCTCGGGCTATTCTTTTGATTTATAAGGGATTTTGCCGATTTCGGAACCACC
ATCAAACAGGATTTTCGCCTGCTGGGGCAAACCAGCGTGGACCGCTTGCTGCAACTCTCTCAGGGCCAGGCGGTGAAGGGCAATCAGCT
GTTGCCCGTCTCACTGGTGAAAAGAAAAACCACCCTGGCGCCCAATACGCAAACCGCCTCTCCCCGCGCGTTGGCCGATTCATTAATGC
AGCTGGCACGACAGGTTTCCCGACTGGAAAGCGGGCAGTGAGCGCAACGCAATTAATGTGAGTTAGCTCACTCATTAGGCACCCCAGGC
TTTACACTTTATGCTTCCGGCTCGTATGTTGTGTGGAATTGTGAGCGGATAACAATTTCACACAGGAAACAGCTATGACCATGATTACG
AATTCGAGCTCGGTACCCGGGGATCCTCTAGAGTCGACCTGCAGGCATGCAAGCTTGGCACTGGCCGTCGTTTTACAACGTCGTGACTG
GGAAAACCCTGGCGTTACCCAACTTAATCGCCTTGCAGCACATCCCCCTTTCGCCAGCTGGCGTAATAGCGAAGAGGCCCGCACCGATC
GCCCTTCCCAACAGTTGCGCAGCCTGAATGGCGAATGGCGCTTTGCCTGGTTTCCGGCACCAGAAGCGGTGCCGGAAAGCTGGCTGGAG
TGCGATCTTCCTGAGGCCGATACTGTCGTCGTCCCCTCAAACTGGCAGATGCACGGTTACGATGCGCCCATCTACACCAACGTGACCTA
TCCCATTACGGTCAATCCGCCGTTTGTTCCCACGGAGAATCCGACGGGTTGTTACTCGCTCACATTTAATGTTGATGAAAGCTGGCTAC
AGGAAGGCCAGACGCGAATTATTTTTGATGGCGTTCCTATTGGTTAAAAAATGAGCTGATTTAACAAAAATTTAATGCGAATTTTAACA
AAATATTAACGTTTACAATTTAAATATTTGCTTATACAATCTTCCTGTTTTTGGGGCTTTTCTGATTATCAACCGGGGTACATATGATT
GACATGCTAGTTTTACGATTACCGTTCATCGATTCTCTTGTTTGCTCCAGACTCTCAGGCAATGACCTGATAGCCTTTGTAGATCTCTC
AAAAATAGCTACCCTCTCCGGCATTAATTTATCAGCTAGAACGGTTGAATATCATATTGATGGTGATTTGACTGTCTCCGGCCTTTCTC
ACCCTTTTGAATCTTTACCTACACATTACTCAGGCATTGCATTTAAAATATATGAGGGTTCTAAAAATTTTTATCCTTGCGTTGAAATA
AAGGCTTCTCCCGCAAAAGTATTACAGGGTCATAATGTTTTTGGTACAACCGATTTAGCTTTATGCTCTGAGGCTTTATTGCTTAATTT
TGCTAATTCTTTGCCTTGCCTGTATGATTTATTGGATGTT

### Supplementary Table 2: Staple sequences

| ID | Description | Sequence |
|---|---|---|
| 1 | Core | AAAAACCAAACCCTCGTTGTGAATATGGTTTGGTC |
| 2 | Core | GGAAGAAGTGTAGCGGTCACGTTATAATCAGCAGACTGATAG |
| 3 | Core | TACGATATAGATAATCGAACAACA |
| 4 | Core | CTTTTGCTTAAGCAATAAAGCGAGTAGA |
| 5 | Core | GTCTGAAATAACATCGGTACGGCCGCGCACGG |
| 6 | Core | GGAAGAGCCAAACAGCTTGCAGGGAACCTAA |
| 7 | Core | AAAATCACCGGAAGCAAACTCTGTAGCT |
| 8 | Core | CCTACATGAAGAACTAAAGGGCAGGGCGGAGCCCCGGGC |
| 9 | Core | CATGTAAAAAGGTAAAGTAATAAGAACG |
| 10 | Core | ATTAAATCAGGTCATTGCCTGTCTAGCTGATAAATTGTAATA |
| 11 | Core | ATAGTCGTCTTTTGCGGTAATGCC |
| 12 | Core | AGTCATGGTCATAGCTGAACTCACTGCCAGT |
| 13 | Core | AACTATTGACGGAAATTTGAGGGAATATAAA |
| 14 | Core | ATCGCGTCTGGAAGTTTCATTCCATATAGAAAGACCATC |
| 15 | Core | AAATATTGAACGGTAATCGTAGCCGGAGACAGTCATAAAAAT |
| 16 | Core | GTCTTTACAGGATTAGTATTCTAACGAGCATAGAACGC |
| 17 | Core | GCACCGCGACGACGCTAATGAACAGCTG |
| 18 | Core | AACTTCATTTTAGAATCGCAAATC |
| 19 | Core | CGTAGAGTCTTTGTTAAGGCCTTCGTTTTCCTACCGAG |
| 20 | Core | CCAATCAAAGGCTTATCCGGTTGCTATT |
| 21 | Core | AGAGGCGATATAATCCTGATTCATCATA |
| 22 | Core | CCGTAATCCCTGAATAATAACGGAATACTACG |
| 23 | Core | AAATGGTATACAGGGCAAGGAAATC |
| 24 | Core | TCCTCATCGTAACCAAGACCGACA |
| 25 | Core | CATTATCTGGCTTTAGGGAATTATGTTTGGATTAC |

| | | |
|---|---|---|
| 26 | Core | ACCCGCCCAATCATTCCTCTGTCC |
| 27 | Core | CGACCAGTCACGCAGCCACCGCTGGCAAAGCGAAAGAAC |
| 28 | Core | CTAAAGGCGTACTATGGTTGCAACAGGAGAGA |
| 29 | Core | TTGGCAGGCAATACAGTGTTTCTGCGCGGGCG |
| 30 | Core | TATACAGGAAATAAAGAAATTTTGCCCGAACGTTAAGACTTT |
| 31 | Core | AAGTATAGTATAAACAGTTAACTGAATTTACCGTTGAGCCAC |
| 32 | Core | ACATTCAGATAGCGTCCAATATTCAGAA |
| 33 | Core | AAACATCTTTACCCTCACCAGTAAAGTGCCCGCCC |
| 34 | Core | GAGATGACCCTAATGCCAGGCTATTTTT |
| 35 | Core | TCCTGAATTTTTTGTTTAACGATCAGAGCGGA |
| 36 | Core | GCCGAAAAATCTAAAGCCAATCAAGGAAATA |
| 37 | Core | AGCGTAGCGCGTTTTCACAAAATCTATGTTAGCAAACGAACGCAACAAA |
| 38 | Core | ACCAATCGATTAAATTGCGCCATTATTA |
| 39 | Core | ATCTTACTTATTTTCAGCGCCGACAGGATTCA |
| 40 | Core | CCCTAAAAGAACCCAGTCACA |
| 41 | Core | GGAAGGGCGAAAATCGGGTTTTTCGCGTTGCTCGT |
| 42 | Core | CAGACCGGAAGCCGCCATTTTGATGGGGTCAGTAC |
| 43 | Core | TAATATTGGAGCAAACAAGAGATCAATATGATATTGCCTTTA |
| 44 | Core | TTCCTTATAGCAAGCAAATCAAATTTTA |
| 45 | Core | ACTACGAGGAGATTTTTTCACGTTGAAACTTGCTTT |
| 46 | Core | AAACAGGCATGTCAATCATATAGATTCAAAAGGGTTATATTT |
| 47 | Core | AACAGGCACCAGTTAAAGGCCGCTTTGTGAATTTCTTA |
| 48 | Core | TTCCTGAGTTATCTAAAATATTCAGTTGTTCAAATAGCAG |
| 49 | Core | AAAGAAACAAGAGAAGATCCGGCT |
| 50 | Core | TTGAGGGTTCTGGTCAGGCTGTATAAGC |
| 51 | Core | TTTAACCGTCAATAGTGAATTCAAAAGAAGATGATATCGCGC |
| 52 | Core | ACGAGCGCCCAATCCAAATAAAATTGAGCACC |
| 53 | Core | AATAAGTCGAAGCCCAATAATTATTTATTCTT |
| 54 | Core | ACGAAATATCATAGATTAAGAAACAATGGAACTGA |
| 55 | Core | TTTCATAGTTGTACCGTAACACTGGGGTTTT |
| 56 | Core | AGGAGCGAGCACTAACAACTAAAACCCTATCACCTAACAGTG |
| 57 | Core | CAAAGTATTAATTAGCGAGTTTCGCCACAGAACGA |
| 58 | Core | TGGGGAGCTATTTGACGACTAAATACCATCAGTTT |
| 59 | Core | ATAACGCAATAGTAAAATGTTTAAATCA |
| 60 | Core | ACGAATCAACCTTCATCTTATACCGAGG |
| 61 | Core | TAATGGTTTGAAATACGCCAA |
| 62 | Core | CGGAACAAGAGCCGTCAATAGGCACAGACAATATCCTCAATC |
| 63 | Core | ATTAAAGGTGAATTATCAAAGGGCACCACGG |
| 64 | Core | GGCAACCCATAGCGTAAGCAGCGACCATTAA |
| 65 | Core | AGAAACGTAAGCAGCCACAAGGAAACGATCTT |
| 66 | Core | AGAGGTCTTTAGGGGGTCAAAAGGCAGT |
| 67 | Core | GGGGACTTTTTCATGAGGACCTGCGAGAATAGAAAGGAGGAT |
| 68 | Core | TTTTAGAACATCCAATAAATCCAATAAC |
| 69 | Core | AAATGTGGTAGATGGCCCGCTTGGGCGC |
| 70 | Core | ACGGATCGTCACCCTCACGATCTAGAATTTT |
| 71 | Core | CGCCATAAGACGACGACAATAGCTGTCT |
| 72 | Core | GCGTATTAGTCTTTAATCGTAAGAATTTACA |
| 73 | Core | AGAGAACGTGAATCAAATGCGTATTTCCAGTCCCC |
| 74 | Core | AACGAAAAAGCGCGAAAAAAAGGCTCCAAAAGG |

| | | |
|---|---|---|
| 75 | Core | TAATTTAGAACGCGAGGCGTTAAGCCTT |
| 76 | Core | ACCAGGCGTGCATCATTAATTTTTTCAC |
| 77 | Core | CAGCCTGACGACAGATGTCGCCTGAAAT |
| 78 | Core | ATTAGTCAGATTGCAAAGTAAGAGTTAAGAAGAGT |
| 79 | Core | CTCGAATGCTCACTGGCGCAT |
| 80 | Core | GGGCAGTCACGACGTTGAATAATTAACAACC |
| 81 | Core | TAAAAACAGGGGTTTTGTTAGCGAATAATATAATAGAT |
| 82 | Core | TCAACCCTCAGCGCCGAATATATTAAGAATA |
| 83 | Core | ATTATACGTGATAATACACATTATCATATCAGAGA |
| 84 | Core | GCAAATCTGCAACAGGAAAAATTGC |
| 85 | Core | ATAATTACTAGAAATTCTTAC |
| 86 | Core | TATCACCGTGCCTTGAGTAACGCGTCATACATGGCCCCTCAG |
| 87 | Core | AAGTAGGGTTAACGCGCTGCCAGCTGCA |
| 88 | Core | CCAGTAGTTAAGCCCTTTTTAAGAAAAGCAAA |
| 89 | Core | TGGCGAAGTTGGGACTTTCCG |
| 90 | Core | CAGTGAGTGATGGTGGTTCCGAAAACCGTCTATCACGATTTA |
| 91 | Core | AAATCAAAGAGAATAACATAACTGAACACAGT |
| 92 | Core | CTGTATGACAACTAGTGTCGA |
| 93 | Core | ATCATAAATAGCGAGAGGCTTAGCAAAGCGGATTGTTCAAAT |
| 94 | Core | TTGAGTAATTTGAGGATTTAGCTGAAAGGCGCGAAAGATAAA |
| 95 | Core | ATAAGAATAAACACCGCTCAA |
| 96 | Core | CGTTGTAATTCACCTTCTGACAAGTATTTTAA |
| 97 | Core | AACCGCCTCATAATTCGGCATAGCAGCA |
| 98 | Core | AAATAGGTCACGTTGGTAGCGAGTCGCGTCTAATTCGC |
| 99 | Core | CAGTATAGCCTGTTTATCAACCCCATCC |
| 100 | Core | TTGCACCTGAAAATAGCAGCCAGAGGGTCATCGATTTTCGGT |
| 101 | Core | CGTCGGAAATGGGACCTGTCGGGGGAGA |
| 102 | Core | AAGAAACTAGAAGATTGCGCAACTAGGG |
| 103 | Core | CCAGAACCTGGCTCATTATACAATTACG |
| 104 | Core | ACGGGTAATAAATTAAGGAATTGCGAATAGTA |
| 105 | Core | CCACGCTGGCCGATTCAAACTATCGGCCCGCT |
| 106 | Core | GCCTTCACCGAAAGCCTCCGCTCACGCCAGC |
| 107 | Core | CAGCATTAAAGACAACCGTCAAAAATCA |
| 108 | Core | ACATCGGAAATTATTTGCACGTAAAAGT |
| 109 | Core | CAACGGTCGCTGAGGCTTGATACCTATCGGTTTATCAGATCT |
| 110 | Core | AAATCGTACAGTACATAAATCAGATGAA |
| 111 | Core | TTAACACACAGGAACACTTGCCTGAGTATTTG |
| 112 | Core | AGGCATAAGAAGTTTTGCCAGACCCTGA |
| 113 | Core | GACGACATTCACCAGAGATTAAAGCCTATTAACCA |
| 114 | Core | AGCTGCTCGTTAATAAAACGAGAATACC |
| 115 | Core | CTTAGAGTACCTTTTAAACAGCTGCGGAGATTTAGACTA |
| 116 | Core | CACCCTCTAATTAGCGTTTGCTACATAC |
| 117 | Core | GAACCGAAAATTGGGCTTGAGTACCTTATGCGATTCAACACT |
| 118 | Core | GCAAGGCAGATAACATAGCCGAACAAAGTGGCAACGGGA |
| 119 | Core | ATGAAACAATTGAGAAGGAAACCGAGGATAGA |
| 120 | Core | GGATGTGAAATTGTTATGGGGTGCACAGTAT |
| 121 | Core | GGCTTGCGACGTTGGGAAGAACAGATAC |
| 122 | Core | TAAATGCCTACTAATAGTAGTTTTCATT |
| 123 | Core | TGCCGTCTGCCTATTTCGGAACCAGAATGGAAAGCCCACCAGAAC |

| | | |
|---|---|---|
| 124 | Core | TGACCATAGCAAAAGGGAGAACAAC |
| 125 | Core | CGAGCCAGACGTTAATAATTTGTATCA |
| 126 | Core | GCTCAGTTTCTGAAACATGAAACAAATAAATCCTCCCGCCGC |
| 127 | Core | AGACGCTACATCAAGAAAACACTTTGAA |
| 128 | Core | AGTACTGACCAATCCGCGAAGTTTAAGACAG |
| 129 | Core | GATTCCTGTTACGGGCAGTGAGCTTTTCCTGTGTGCTG |
| 130 | Core | GGTATTAAGGAATCATTACCGAACGCTA |
| 131 | Core | GTTCATCAAATAAAACGCGACTCTAGAGGATCGGG |
| 132 | Core | AGCCTTTAATTGGATAGTTGAACCGCCACCCTCATAGGTG |
| 133 | Core | ACAGAGGCCTGAGATTCTTTGATTAGTAATGG |
| 134 | Core | AACGAGATCAGGATTAGAGAGCTTAATT |
| 135 | Core | TACCAAGTTATACTTCTGAATCACCAGA |
| 136 | Core | CAGTAGGTGTTCAGCTAATGCGTAGAAA |
| 137 | Core | AGGATGACCATAGACTGACTAATGAAATCTACATTCAGCAGGCGCGTAC |
| 138 | Core | TTTCAACCAAGGCAAAGAATTTAGATAC |
| 139 | Core | TTGAAATTAAGATAGCTTAACTAT |
| 140 | Core | CTATTATCGAGCTTCAAAGCGTATGCAA |
| 141 | Core | CAGGGTGCAAAATCCCTTATAGACTCCAACGTCAAAAGCCGG |
| 142 | Core | GAGCTTGTTAATGCGCCGCTAATTTTAGCGCCTGCTGCTGAA |
| 143 | Core | CGAACGTTAACCACCACACCCCCAGAATTGAG |
| 144 | Core | GTGTGATAAATAAGTGAGAAT |
| 145 | Core | GCTATATAGCATTAACCCTCAGAGA |
| 146 | Core | AGGAGAGCCGGCAGTCTTGCCCCCGAGAGGGAGGG |
| 147 | Core | CGGCCTCCAGCCAGAGGGCGAGCCCCAA |
| 148 | Core | CCAAAACAAAATAGGCTGGCTGACGTAACAA |
| 149 | Core | GGCGGTTAGAATAGCCCGAGAAGTCCACTATTAAAAAGGAAG |
| 150 | Core | ATAAAGGTTACCAGCGCTAATTCAAAAACAGC |
| 151 | Core | ATTGCCCCCAGCAGGCGAAAAGGCCCACTACGTGACGGAACC |
| 152 | Core | TTTTAAAACATAACAGTAATGGAACGCTATTAGAACGC |
| 153 | Core | AATTGGGTAACGCCAGGCTGTAGCCAGCTAGTAAACGT |
| 154 | Edge | TTACCCAGAACAACATTATTACAGGTTTTTTTTTTTTTTTT |
| 155 | Edge | TTTTTTTTTTTTTTTTAATAAGAGAATA |
| 156 | Edge | TTTTTTTTTTTTTTTTCCAGTTTGGGAGCGGGCTTTTTTTTTTTTTTT |
| 157 | Edge | GGTTGAGGCAGGTCAGTTTTTTTTTTTTTTT |
| 158 | Edge | TTTTTTTTTTTTTTTGATTAAGACTCCTTATCCAAAAGGAAT |
| 159 | Edge | TTTTTTTTTTTTTTTTCTTCGCTATTACAATT |
| 160 | Edge | TTTTTTTTTTTTTTTCTTGCGGGAGAAGCGCATTTTTTTTTTTTTTTT |
| 161 | Edge | TTTTTTTTTTTTTTGGGAATTAGAGAAACAATGAATTTTTTTTTTTTTTT |
| 162 | Edge | TCAGACTGACAGAATCAAGTTTGTTTTTTTTTTTTTTT |
| 163 | Edge | TTTTTTTTTTTTTTTGGTCGAGGTGCCGTAAAGCAGCACGT |
| 164 | Edge | TTTTTTTTTTTTTTTTTTAATCATTTACCAGACTTTTTTTTTTTTTTT |
| 165 | Edge | TTTTTTTTTTTTTCATTCTGGCCAAATTCGACAACTCTTTTTTTTTTTTT |
| 166 | Edge | TTTTTTTTTTTTTTTACCGGATATTCA |
| 167 | Edge | TTTTTTTTTTTTTTTTAGACGGGAAACTGGCATTTTTTTTTTTTTTTT |
| 168 | Edge | TTTTTTTTTTTTTTTCAGCAAGCGGTCCACGCTGCCCAAAT |
| 169 | Edge | CTGAGAGAGTTGTTTTTTTTTTTTTTT |
| 170 | Edge | CAATGACAACAACCATTTTTTTTTTTTTTTT |
| 171 | Edge | TTTTTTTTTTTTTTTTGAGAGATCTACAAGGAGAGG |
| 172 | Edge | TCACCAGTACAAACTATTTTTTTTTTTTTTT |

| | | |
|---|---|---|
| 173 | Edge | TTTTTTTTTTTTTGGCAATTCATCAAATTATTCATTTTTTTTTTTTTTTT |
| 174 | Edge | TAAAGTTACCGCACTCATCGAGAACTTTTTTTTTTTTTTT |
| 175 | Edge | TTTTTTTTTTTTTTTCACCCTCAGAACCGCC |
| 176 | Edge | TTTTTTTTTTTTTAGGTTTAACGTCAATATATGTGAGTTTTTTTTTTTTT |
| 177 | Edge | CCACACAACATACGTTTTTTTTTTTTT |
| 178 | Edge | TTTTTTTTTTTTTTTGCTAGGGCGAGTAAAAGATTTTTTTTTTTTTTT |
| 179 | Edge | TTTTTTTTTTTTTTTAGTTGATTCCCAATTCTGCGAACCTCA |
| 180 | Edge | TTATTTAGAGCCTAATTTGCCAGTTTTTTTTTTTTTTTTT |
| 181 | Edge | TTTTTTTTTTTTTTTACGGCGGAT |
| 182 | Edge | TTTTTTTTTTTTTTTATATGCGTTAAGTCCTGATTTTTTTTTTTTTTT |
| 183 | Edge | TTTTTTTTTTTTTTTACGATTGGCCTTGATA |
| 184 | Edge | TTTTTTTTTTTTTTTCAACGCCTGTAGCATT |
| 185 | Edge | TTTTTTTTTTTTTTTGGCTTTGAGCCGGAACGATTTTTTTTTTTTTTT |
| 186 | Edge | TTTTTTTTTTTTTTTAAGCAAGCCGTTT |
| 187 | Edge | TTTTTTTTTTTTTATGTGTAGGTAAGTACCCCGGTTGTTTTTTTTTTTTT |
| 188 | Edge | ATCGTCATAAATATTCATTTTTTTTTTTTTTTTT |
| 189 | Edge | TTTTTTTTTTTTTTTGTTAATTTCATCT |
| 190 | Edge | TTTTTTTTTTTTTGTATTAAATCCTGCGTAGATTTCTTTTTTTTTTTTTT |
| 191 | Edge | GCCATATAAGAGCAAGCCAGCCCGACTTGAGCCATGGTT |
| 192 | Edge | GTAGCTAGTACCAAAAACATTCATAAAGCTAAATCGGTTTTTTTTTTTTT |
| 193 | Edge | ATAACGTGCTTTTTTTTTTTTTTTTTT |
| 194 | Edge | TTTTTTTTTTTTTTTAAAATACCGAACGAACCACCAGTGAGAATTAAC |
| 195 | Edge | TTTTTTTTTTTTTTTACAAAATAAACA |
| 196 | Edge | TTTTTTTTTTTTTTTACAAGAAAAACCTCCCGATTTTTTTTTTTTTTT |
| 197 | Edge | TTTTTTTTTTTTTTTGACGATAAAAAGATTAAGTTTTTTTTTTTTTTT |
| 198 | Edge | TTTTTTTTTTTTTCAATTACCTGAGTATCAAAATCATTTTTTTTTTTTTT |
| 199 | Edge | GGTACGGCCAGTGCCAAGCTTTTTTTTTTTTTTT |
| 200 | Edge | TTTTTTTTTTTTTTGAATAACCTTGAAATATATTTATTTTTTTTTTTTTT |
| 201 | Edge | CACTAAAACACTTTTTTTTTTTTTTTT |
| 202 | Edge | TTTTTTTTTTTTTTTAACCAATATGGGAACAATTTTTTTTTTTTTTTT |
| 203 | Edge | TACGTCACAATCAATAGAATTTTTTTTTTTTTTT |
| 204 | Edge | TTTTTTTTTTTTTTTAGAAAGATTCATCAGTTGA |
| 205 | Edge | TTTTTTTTTTTTTGTGGCATCAATTAATGCCTGAGTATTTTTTTTTTTTT |
| 206 | Edge | TTTTTTTTTTTTTTTTTGCATGCCTGCATTAATTTTTTTTTTTTTTTT |
| 207 | Edge | CCAGCGAAAGAGTAATCTTGACAAGATTTTTTTTTTTTTTT |
| 208 | Edge | TTTTTTTTTTTTTTTGAATCCCCCTCAAATGCTT |
| 209 | Edge | AGAGGCTGAGACTCCTTTTTTTTTTTTTTTT |
| 210 | Edge | ACAAACACAGAGATACATCGCCATTATTTTTTTTTTTTTTT |
| 211 | Edge | TTTTTTTTTTTTTTTCAAGAGAAGGATTAGG |
| 212 | Edge | TTTTTTTTTTTTTGAATTGAGGAAGTTATCAGATGATTTTTTTTTTTTTT |
| 213 | Edge | CAGAACAATATTTTTTTTTTTTTTTTT |
| 214 | Edge | TTTTTTTTTTTTTAGCCGGAAGCATAAAGTGTCCTGGCC |
| 215 | Edge | TGACCGTTTCTCCGGGAACGCAAATCAGCTCATTTTTTTTTTTTTTTTTT |
| 216 | Edge | TTTTTTTTTTTTTTTGGTAATAAGTTTTAAC |
| 217 | Edge | TTTTTTTTTTTTTTGTCTGTCCATAATAAAAGGGATTTTTTTTTTTTTTT |
| 218 | Edge | TTTTTTTTTTTTTTTCCTCGTTAGAATCAGAGCGTAATATC |
| 219 | Edge | AATTGCTCCTTTTGATAAGTTTTTTTTTTTTTTT |
| 220 | Edge | CATCGGACAGCCCTGCTAAACAACTTTCAACAGTTTTTTTTTTTTTTT |
| 221 | Edge | TTTTTTTTTTTTTTTAACCGCCTCCCTCAGACCAGAGC |

| | | |
|---|---|---|
| 222 | Edge | TCTGACAGAGGCATTTTCGAGCCAGTTTTTTTTTTTTTTT |
| 223 | Edge | TTTTTTTTTTTTTTTTTTCAGCGGAGTTCCATGTCATAAGG |
| 224 | Edge | TTTTTTTTTTTTTTTCGCCCACGCATAACCG |
| 225 | Edge | AATTACTTAGGACTAAATAGCAACGGCTACAGATTTTTTTTTTTTTTT |
| 226 | Edge | CAAGTTTTTTGGTTTTTTTTTTTTTTT |
| 227 | Edge | TTTTTTTTTTTTTTTCCTTTAGCGCACCACCGGTTTTTTTTTTTTTTT |
| 228 | Edge | TTTTTTTTTTTTTTTGAATCGGCCGAGTGTTGTTTTTTTTTTTTTTTT |
| 229 | Edge | TTTTTTTTTTTTTCATCTTTGACCC |
| 230 | Edge | TTTTTTTTTTTTTATAATCAGAAAATCGGTGCGGGCCTTTTTTTTTTTTT |
| 231 | Edge | GATACAGGAGTGTACTTTTTTTTTTTTTTTT |
| 232 | Edge | TTTTTTTTTTTTTTTGGCGCAGACAATTTCAACTTTTTTTTTTTTTTT |
| 233 | Edge | GGAGGTTTAGTACCGCTTTTTTTTTTTTTTT |
| 234 | Edge | TTTTTTTTTTTTTACCGCCAGCCATAACAGTTGAAAGTTTTTTTTTTTTT |
| 235 | Edge | TTTTTTTTTTTTTTTATAGCAATAGCT |
| 236 | Handles | AATAAGTTTTGCAAGCCCAATAGGGGATAAGTTGTGCTACTCCAGTTC |
| 237 | Handles | ACATAGCTTACATTTAACAATAATAACGTTGTGCTACTCCAGTTC |
| 238 | Handles | CCTTTTTGAATGGCGTCAGTATTGTGCTACTCCAGTTC |
| 239 | Handles | CGTAACCAATTCATCAACATTTTGTGCTACTCCAGTTC |
| 240 | Handles | CACCAACCGATATTCATTACCATTATTGTGCTACTCCAGTTC |
| 241 | Handles | CCACCCTCATTTTCTTGATATTTGTGCTACTCCAGTTC |
| 242 | Handles | AACTTTGAAAGAGGAGAAACATTGTGCTACTCCAGTTC |
| 243 | Handles | CAAGGCGCGCCATTGCCGGAATTGTGCTACTCCAGTTC |
| 244 | Handles | CATAGCCCCCTTAAGTCACCATTGTGCTACTCCAGTTC |
| 245 | Handles | TTTCCCTGAATTACCTTTTTTACCTTTTTTGTGCTACTCCAGTTC |
| 246 | Handles | AACGGTGTACAGACTGAATAATTGTGCTACTCCAGTTC |
| 247 | Handles | GATTCGCGGGTTAGAACCTACCATTTTGTTGTGCTACTCCAGTTC |
| 248 | Guides | AGAGTAGGATTTCGCCAACATGTTTTAAAAACC |
| 249 | Guides | ACGGTGACCTGTTTAGCTGAATATAATGCCAAC |
| 250 | Guides | CGTAGCAATTTAGTTCTAAAGTACGGTGTTTTA |
| 251 | Guides | GCTTAATGCGTTAAATGTAAATGCTGATCTTGAAATGAGCGTT |
| 252 | Guides | AAGCCAACGGAATCTAGGTTGGGTTATATAGATTAAGCAACTG |
| 253 | Guides | TTTAACAACCGACCCAATCGCAAGACAAAATTAATCTCACTGC |
| 254 | Guides | TTTAGGCCTAAATTGAGAAAACTTTTTCCTTCTGTTCCTAGAT |
| 255 | Guides Removal | GGTTTTTAAAACATGTTGGCGAAATCCTACTCT |
| 256 | Guides Removal | GTTGGCATTATATTCAGCTAAACAGGTCACCGT |
| 257 | Guides Removal | TAAAACACCGTACTTTAGAACTAAATTGCTACG |
| 258 | Guides Removal | AACGCTCATTTCAAGATCAGCATTTACATTTAACGCATTAAGC |
| 259 | Guides Removal | CAGTTGCTTAATCTATATAACCCAACCTAGATTCCGTTGGCTT |
| 260 | Guides Removal | GCAGTGAGATTAATTTTGTCTTGCGATTGGGTCGGTTGTTAAA |
| 261 | Guides Removal | ATCTAGGAACAGAAGGAAAAAGTTTTCTCAATTTAGGCCTAAA |

To fold the robots, scaffold and staple DNA were mixed at a ratio of 1:10, respectively, in Tris-Acetate-EDTA buffer supplemented with 10 mM $MgCl_2$. The mixture was subjected to a temperature-annealing ramp in the following sequence: 1) from 85°C to 60° C, 5 min/°C; 2) from 60 °C to 25 °C, 75 min/°C. Subsequently, excess staples were removed by centrifugal filtration using Amicon Ultra-0.5mL 100K MWCO centrifugal filters (Millipore).

**Payload synthesis**

GFP was fused to loading-sequence DNA (5AmMC6/GAACTGGAGTAGCAC Integrated DNA Technologies) by EDC conjugation according to the manufacturer's instructions. Anti-human p75/AIRM Fab' fragments were obtained by digesting whole IgG using a Fab' generation kit (Pierce) according to the manufacturer's instructions. After purification, Fab' fragments were fused to loading sequence DNA by EDC conjugation.

**Robot loading and purification**

100 pmol folded robots were loaded with autoinducer and payload (at a 3:1 ratio) by incubation at a 5-fold molar excess of mixture to loading sites. Loading was performed for 2 hours on a rotary shaker at room temperature in folding buffer (10 mM $MgCl_2$ in 1X TAE). Finally, loaded robots were cleaned by centrifugal filtration with a 100K MWCO Amicon column (Millipore) as described above.

Loading in this design was done stochastically. However, by redesigning the loading site sequences and autoinducer/payload specificities, loading can be directed to specific sites. However, stochastic loading was effective (albeit potentially less optimal than directed loading), for the following reasons: a) robots containing only autoinducer can serve as autoinducer sources indicating population size; b) robots containing only payload (GFP/Fab') respond to external autoinducer and contribute to the readout; and c) robots containing both serve both functions.

**Supplementary Note 2: QS system design**

5'-amine-modified linker oligonucleotide (5AmMC6/TTTTTGAACTGGAGTAGCAC, Integrated DNA Technologies) was conjugated using the heterobifunctional crosslinker SMCC to the C-terminal thiol group in Lys-Pro-Leu-Gly-Met-Trp-Ser-Arg-Cys (custom ordered from American Peptide Company), containing the cleavage site of MMP-2. according to the manufacturer instruction, at a DNA:peptide ratio of 1:2. After quenching with 2-mercaptoethanol and purification, the oligonucleotide-peptide hybrid was further conjugated with PDGF using EDC crosslinking, purified and verified with spectrophotometry to yield to complete autoinducer. The cleaved autoinducer maintained its ability to bind to anti-PDGF antibodies as well as to the PDGF aptamer.

Kinetics of peptide cleavage by MMP-2 was measured by fluorometry using a fluorogenic MMP-2/MMP-2 substrate (5 μM) and human recombinant MMP-2 in assay buffer (Tris-EDTA containing 150 mM NaCl, 10 mM $MgCl_2$ and 1 μM $ZnSO_4$, pH 7.5) at room temperature. The desired concentration of MMP-2 for this study was fixed at 5 μg/mL **(Fig. S1)**.

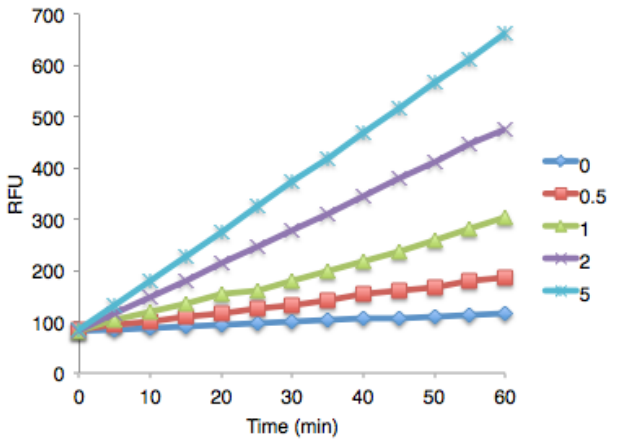


**Fig. S1:** MMP-2 calibration assay, used to determine desired MMP-2 concentration for the purpose of activating QS in robots for this study (see above for detail).

To evaluate the kinetics of autoinducer release from robots, autoinducer-loaded robots were exposed to MMP-2 (5 μg/mL) for 1 h at room temperature, after which the samples were measured directly in a PDGF ELISA **(Fig. S2).**

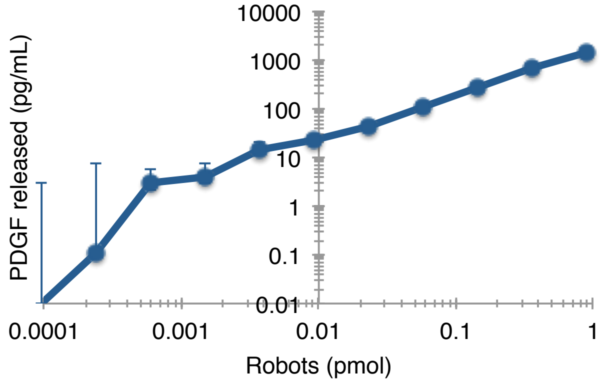


**Fig. S2: Autoinducer release from MMP-2 treated robots.**

**Supplementary Note 3: Cell culture**

Jurkat cells were obtained from American Type Culture Collection (ATCC) and maintained at 37 deg. and 5% $CO_2$ in RPMI 1640 containing 10% fetal calf serum. Prior to incubation with robots, cells were diluted to a density of 100,000 cells/mL in 96 well plates and activated with 200 ng/mL of recombinant human IL-6 (Peprotech) overnight.
Following activation, the cells were treated with varying concentrations of either free anti-p75/AIRM Fab' fragments (cross-linked by 25 ug/mL secondary anti-mouse IgM), or the equivalent amount of Fab' fragments loaded into QS-regulated robots, for 24 hours. Following this period, the cells were analyzed for cell cycle distribution using propidium iodide as previously described.

**Supplementary Note 4: Dynamic light scattering and flow cytometry**

Dynamic light scattering was performed using a Malvern Zetasizer Nano instrument using various concentrations of robots in Tris-EDTA buffer supplemented with 10 mM $MgCl_2$. The minimal robot concentration that enabled reliable detection (based on good correlation function) was 29 pM, and the results obtained were good as a qualitative confirmation of QS-driven switch from closed to open state.
The advantage of flow cytometry is the use of target-coated microspheres, which isolate from any population only the robots that open and directly bind them, allowing much more reliable measurements and at lower population densities. Flow cytometry was performed using an Accuri C6 flow cytometer equipped with 488 nm and 640 nm lasers, and analyzed with FlowPlus software. Cell cycle analysis was done using propidium iodide as previously described.
